# *Drosophila melanogaster Peroxisome Biogenesis Factor 7* plays a role in embryonic development

**DOI:** 10.1101/757815

**Authors:** C Pridie, AJ Simmonds

**Affiliations:** Department of Cell Biology, Faculty of Medicine and Dentistry, University of Alberta, Edmonton, Alberta T6G 2H7, Canada

**Keywords:** peroxisome, Peroxin, *Drosophila melanogaster*, embryogenesis, central nervous system, forward genetic screen, TRiP-KO, Gal4-UAS

## Abstract

Peroxisomes are organelles responsible for aspects of lipid metabolism and management of reactive oxygen species. Peroxisome Biogenesis Factor (*Peroxin, Pex*) genes encode proteins essential to peroxisome biogenesis or function. In yeast and mammals, PEROXIN7 acts as a cytosolic receptor protein that targets a subset of enzymes for peroxisome matrix import.

Proteins targeted by PEROXIN7 contain a peroxisome targeting sequence 2 (PTS2) motif. The PTS2 was not found in the *D. melanogaster* homologs of proteins that are PEROXIN7 targets in yeast or mammals, however comparative genomics suggest a *Pex7* homolog is present in the *D. melanogaster* genome. Herein we report novel, tissue-specific patterns for transcription and translation of *Pex7* in the *D. melanogaster* embryo that appear to be strongest in presumptive neuronal lineages. We also show that targeted somatic *Pex7* knockout in neural precursors via targeted somatic CRISPR knockout affected survival of mutant embryos. *Pex7* over-expression via Gal4-UAS also reduced adult survival but was not deleterious at the embryo stage. Notably, targeted somatic rescue of *Pex7* in the neural precursors of *Pex7* homozygous mutants also impaired embryo survival. We conclude that *D. melanogaster* has tissue-specific developmental requirements of *Pex7* expression. This may be related to the requirement for peroxisome-mediated lipid synthesis in cells of the central nervous system.

## Introduction

Peroxisomes are single-membrane-bound organelles found in most eukaryotes. Their role varies by organism with a core functionality of non-energetic fatty acid β-oxidation and detoxification of reactive oxygen species. In fungi and plants peroxisomes are wholly responsible for fatty acid oxidation, whereas in animals this occurs mostly in mitochondria. In mammals peroxisomes are the site of bile acid and ether lipid (phospholipid) synthesis, the latter being integral to the plasma membranes of white matter neurons (Platta and Erdmann, 2007). Peroxisome biogenesis occurs by two mechanisms, fission of extant mature peroxisomes and budding of new peroxisomes from the ER (Agrawal *et al*., 2016; Yuan *et al*., 2015). The biogenesis and function of peroxisomes, e.g. client enzyme import and membrane assembly, requires proteins called Peroxisome Biogenesis Factors (Peroxin, Pex). Yeast, plants and mammals employ two cytosolic receptors to mediate import, Pex5 and Pex7. Pex5 recognizes its clients by the conserved C-terminal tripeptide (S/A/C)-(K/R/L)-(L/M), termed Peroxisome Targeting Signal 1 (PTS1). Pex7, together with a species-specific accessory protein, recognizes the variable N-terminal sequence (R/K)-(L/V/I)-XXXXX-(H/Q)-(L/M), X being any residue, termed PTS2. The Pex7 binding partner includes Pex5 in plants and fungi, the long isoform Pex5L in mammals or Pex18/21 in *S. cerevisiae* (Montilla-Martinez *et al*., 2015; Cross *et al*., 2015; Platta *et al*., 2015, Emmanouilidis *et al*., 2015). In terms of protein structure, sequence analysis suggests Pex7 contains a series of six or seven WD40 domain repeats, whose canonical tertiary structure is a β-propeller.

Accordingly, Pex7 is proposed to act as a platform for transient protein interactions as has been observed for Pex7-Pex21 complexes in yeast (Pan *et al*., 2013; Emmanouilidis *et al*., 2015). Not all eukaryotes conserve the PTS2 pathway. *C. elegans* and some unicellular diatoms employ only the PTS1 tripeptide and their genome does not include a *Pex7* homologue (Motley *et al*., 2000; Gonzalez *et al*., 2011). The *D. melanogaster* genome encodes a putative *Pex7* homolog only somewhat characterized. To date, different patterns of co-localization of Pex7/peroxisome have been reported, from 0% to 50% co-localized depending on the reporters used (Faust *et al*., 2012; Baron *et al*., 2016). There is a requirement for *Pex7* in *D. melanogaster* lipid processing, as homozygous *Pex7* mutant larvae have higher levels of circulating free fatty acids and are larger than their wild-type counterparts. Mutants also displayed neurological defects as pupae that did not eclose into adults (Di Cara *et al*., 2019). Given the known roles peroxisomes play in lipid processing within the context of the observed effects of *Pex7* loss, a peroxisome-related role for *Pex7* in *Drosophila* may be conjectured, however further characterization is required to elucidate its specific function and determine if *Pex7* is involved in *Drosophila* peroxisome biogenesis.

To explore the role for Pex7 during early fly development, we examined the endogenous spatial pattern of embryonic transcription for several *Drosophila Peroxins*. Using a novel antibody, we then observed *Pex7* encoded a protein present at relatively high levels in a variable subset of cells during embryogenesis. Finally, we discovered *Pex7* mutant phenotypes mirrored some aspects of mutations in two other Peroxins, *Pex1* and *Pex14* (Mast *et al*., 2011; Faust *et al.,* 2012; Baron *et al.,* 2016). Together, these data suggest that *D. melanogaster Pex7* function is required for normal development of a specific subset of cells, likely of neuronal lineage.

## Materials and Methods

### Drosophila stocks, embryo collection and viability assays

1.0-1.5 g of *w^1118^* embryos were collected from 150 mm x 15 mm apple juice/agar plates supplemented with fresh yeast paste, from a 46 cm x 46 cm x 61 cm clear plastic cage deep filled with adult flies. Collections were performed for 2-4 h, as required. The plates were removed from the cage and the embryos aged on the collection plates at 25°C until the desired developmental stage was reached. Aged embryos were washed off the plates in flowing tap water through a three-stage brass sieve, de-chorionated by submersion in 1:1 bleach:water (Clorox) for 90 s, rinsed 3-4 min rinse in flowing tap water and then either flash-frozen in liquid nitrogen for downstream quantitative assays or preserved via paraformaldehyde (4 %) fixation and heptane/methanol extraction, as described previously (Lécuyer *et al.,* 2008), for downstream qualitative assays. Fly stocks used in this study were obtained from the Bloomington *Drosophila* Stock Center, the Kyoto Stock Center (DGRC) or developed *de novo*.

BDSC stocks

*w*^1118^

*w^−^*; *CyO*

*w*^1118^; *sna*^Sco^/CyO, P{ActGFP.w[-]}CC2

*w*\* *y^1^*; Mi{y^+mDint2^=MIC}*Pex7*

P{*w*^+mW.hs^-*elav^C155^*}; P{*y*^+t7.7^ *w*^+mC^-UAS-Cas9.P2}attP40/*CyO*

*w*\*; P{*y*^+t7.7^ *w*^+mC^-UAS-Cas9.P2}attP40/*CyO*;P{w^+mW.hs^-*pnr*^MD237^}/*TM6B*, *Tb*^1^

*y*^1^ *sc*\* *v*^1^; P{*y*^+t7.7^ *v*^+t1.8^-QUAS sgRNA}attP40/*CyO*

*y*^1^ *sc*\* *v*^1^; P{*y*^+t7.7^ *v*^+t1.8^-*Pex7*sgRNA}attP40/*CyO*

*y*^1^ *sc*\* *v*^1^; P{*y*^+t7.7^ *v*^+t1.8^-*Pex14*sgRNA}attP40/*CyO*

*w*\* *y*\*; P{*w*^+mC^-UASp-*Pex7*}attP40

*w*\* *y*\*; P{*w*^+mC^ -UASp-*Pex1*}; *Pex1*\*/*TM6*, *Tb*

Kyoto DGRC stock

*w*^67c23^ *y*^1^; P{*w*^+mC^=GSV6}GS15386/*Sb*^1^, *Ser*^1^, *TM3*

UAS-*Pex1* and UAS-*Pex7* transgenic strains, the latter produced *de novo* for this study, were established by Best Gene Inc., Chino Hills CA, USA (https://www.thebestgene.com) using ΦC31 Integrase-mediated germline integration (Bischof *et al.,* 2007; Bateman *et al.,* 2006). The UAS-*Pex7* cassette was inserted as the plasmid pPMW-attB-*Pex7*, derived from pPMW-attB (Addgene, contributor F Perronet) and pENTR-*Pex7* (Baron *et al*., 2016) using Gateway LR Clonase II (ThermoFisher), then transformed into strain *y*^1^ M{vas-int.Dm}ZH-2A *w*\*; M{3xP3-RFP.attP’}ZH-51C (BDSC).

For the adult survivorship assays, five virgin females were mated to three age-matched males, ±24 h, for 4 d after which the P generation was discarded. Virgin females used were balanced with *CyO* or *CyO*-GFP and/or *Sb*^1^ *Ser*^1^ *TM3* as appropriate. For egg/larva counting assays, ten virgin females were mated to eight age-matched males, ±24 h, for 4 d after which the P generation were discarded. Egg/larva score crosses were housed in small (35 mm diameter) clear embryo collection cages (Genesee Scientific) atop 35 x 10 mm Petri dishes (ThermoFisher) containing apple juice agar medium (CSH Protocols, 2011) supplemented with fresh yeast paste. Dishes were changed after 48 hrs and eggs were counted then the plates were aged 48 h and larvae were counted.

### RNA probe synthesis and *in situ* mRNA analysis

Digoxigenin-labeled anti-sense RNA probes to each *Pex* mRNA were incubated with staged collections of preserved embryos from strain *w*^1118^ using fluorescent *in situ* hybridization (FISH). Plasmids containing cDNA clones of *D. melanogaster Peroxins* (Baron *et al*., 2016), were used as probe templates. Embryo preparation, probe preparation and FISH were all performed as described previously (Lécuyer *et al*., 2008; Wilk *et al*., 2010). Probe synthesis used the Roche DIG RNA Labeling Kit (Millipore-Sigma) and fluorescent signal was generated via tyramide signal amplification using the TSA Individual Cyanine 3 Tyramide Reagent Pack (PerkinElmer), both following manufacturer’s instructions. For embryos over 16 h old, an additional permeabilization step was added between the first post-fixation step and Proteinase K incubation.

An adaptation of a solvent mix published previously (Rand *et al*., 2010), composed of 900 μl (R)-(+)-Limonene (technical grade, Sigma-Aldrich), 50 μl Cocamide DEA (85 % v/v, BiOrigins) and 50 μl Tween-20 (20 % v/v, Sigma-Aldrich), was diluted 1:100 in autoclave-sterilized, 0.22 µm-filtered 1x PBS. Embryos were incubated 10 min in the dilution, washed 3x 5 min in sterile, filtered 1x PBS then the published FISH protocol was followed subsequently.

### Custom antibody generation, purification and *in situ* protein analysis

Two adult rabbits were injected with the chemically-synthesized peptide EQNSNTNSSSTDGQSLGELC, corresponding to predicted *D. melanogaster* Pex7 residues 46-65. Crude serum was isolated from each final bleed by centrifugation and combined then antibodies were precipitated by affinity chromatography using the epitope as bait. Antibody titer was determined by ELISA (Pacific Immunology, Ramona CA). Purified antibody was assayed by western blotting of lysate from *E. coli* cell strain BL21-DE3 expressing glutathione-s-transferase (GST)-Pex7, lysate of Schneider-2 (S2) cells expressing FLAG-Pex7, and lysate of temporally-staged embryos from *D. melanogaster* strain *w^1118^* (Figure 3). GST-Pex7 was expressed by transformed pDEST15-*Pex7* and FLAG-Pex7 was expressed by transfected pAFW-*Pex7*, both sub-cloned from pENTR-*Pex7* (gift of M. Anderson-Baron; Baron *et al.,* 2016).

### Quantitative RT-PCR analysis

Total RNA was isolated from frozen, staged *w^1118^* embryos or adults, as appropriate, via chloroform/phenol-guanidinium isothiocyanate extraction (TRIzol, ThermoFisher) and total cDNA was transcribed using the iScript Select cDNA synthesis kit (BioRad) following manufacturer protocols. Primers were designed using Fly PrimerBank (Hu *et al*., 2013) and verified for target specificity within the fly genome using NCBI Basic Local Alignment Search Tool (BLAST). Primer synthesis was performed by Integrated DNA Technologies (Iowa, USA). Two primer pairs for each gene were validated for amplification efficiency by serial primer dilution (Livak and Schmittgen, 2001).

*Pex7-1*: 5’-ATGCAGACACAGACACACACC, 5’-CAGCAAGTAGTTAGCCTCGAAAG
*Pex7-3*: 5’-CCAACACAAATTCCTCATCCACA, 5’-GTCGAACAATCCATCGGACCA
*Act5C-2*: 5’-AGCGTGAAATCGTCCGTGAC, 5’-GCAAGCCTCCATTCCCAAG
*Act5C-3*: 5’-AGGCCAACCGTGAGAAGATG, 5’-GGGGAAGGGCATAACCCTC

PCR reactions were carried out using SYBR Green PCR Master Mix (Applied Biosystems/ThermoFisher) and black, semi-skirted 96-well twin.tec real-time PCR plates (Eppendorf) in a Mastercycler Realplex^2^ EP gradient S thermal cycler (Eppendorf). Individual reactions totaled 20 μl and used 100 ng raw cDNA as template. The program used was 2 min at 98 °C, 40x (15 s at 98 °C, 60 s at 61.5 °C), followed by a melting curve from 55-98 °C.

### Protein analysis and western blotting

Qualitative and quantitative protein detection were each performed via immunofluorescence and western blotting, respectively. Immunofluorescence was performed as described previously (Lécuyer *et al*., 2008; Wilk *et al*., 2010). Semi-dry western blotting was performed with a Trans-Blot^®^ Turbo transfer system (Bio-Rad) and imaged with an Odyssey infrared imaging scanner (LI-COR-Biosciences). Relative band intensity was quantified with Odyssey V3.0 software (LI-COR-Biosciences). Endogenous *D. melanogaster* Pex7 was detected using the novel Pex7 antibody described above. For immunoflourescent detection of endogenous protein in embryos, an additional post-rehydration permeabilization step was performed for embryos over 16 h using the mixture and protocol noted in the RNA *in situ* section above. Mouse monoclonal primary antibodies α-βTub56D (E7, contributor M Klymkowsky), αElav (9F8A9, contributor GM Rubin), αNrg (BP 104, contributor C Goodman), αRepo (8D12, contributor C Goodman) and αWg (4D4, contributor SM Cohen) were obtained from the Developmental Studies Hybridoma Bank, created by the NICHD of the NIH and maintained at The University of Iowa, Department of Biology, Iowa City, IA 52242. Fluorophore-conjugated reporter antibodies used were DonkeyαMouse-AlexaFluor488 (ThermoFisher), DonkeyαMouse-AlexaFluor690 (ThermoFisher), DonkeyαRabbit-CF555 (Millipore-Sigma) and DonkeyαRabbit-AlexaFluor780 (ThermoFisher). Antibody titer varied by assay with the following dilutions used as baselines:

Immunofluorescence: 1° @ 1:20 of [stock], 2° @ 1:100 of [stock]
Western blotting: 1° @ 1:1000 of [stock], 2° @ 1:5000 of [stock]

### Cell culture care and transfection

Transformation-competent *E. coli*, strains Top10 (ThermoFisher) and BL21 (New England BioLabs), were grown on standard LB+agar with appropriate antibiotic supplement. S2 cells were grown in either HyClone SFX-Insect medium or SFM4-Insect medium plus L-Glutamine (GE Healthcare Life Sciences) supplemented with 100 units/ml penicillin and 100 μg/ml streptomycin (ThermoFisher).

### Microscopy and image analysis

Widefield mages of whole-mount embryos were captured over a total Z-distance of 610 nm with an AxioCam HRm camera (Zeiss) using an x20 apochromatic objective lens (NA = 0.75, Zeiss) on a Zeiss Observer.Z1 microscope controlled by ZEN 2 Blue Edition software (Zeiss). Post-imaging light deconvolution was performed with Huygens Professional software (Scientific Volume Imaging) using X and Y sampling intervals of 189 nm as per the online Scientific Volume Imaging Nyquist Calculator application (https://svi.nl/Nyquist Calculator). Images were rendered as maximum orthogonal projections from de-convolved data using Bitplane Imaris software (Bitplane/Oxford Instruments) and assembled using Adobe Photoshop CS6 software, 64-bit version (Adobe).

### Rescue verification

To verify construct expression in rescue crosses, adult flies of each strain were collected and stored for 1 h in disposable transparent 8 oz. Drosophila bottles (Fisherbrand) then transferred to a fresh bottle and frozen overnight at −80 °C. Molecular verification was performed by PCR amplification of adult gDNA using Phusion High-Fidelity DNA polymerase (ThermoFisher) in 50 µl volumes following manufacturer’s instructions. The targets were a 1.5 kb fragment from the MiMIC transposon’s eGFP CDS, inserted 35-37 bp from the 3’ end of the third exon of the *Pex7* locus (Nagarkar-Jaiswal *et al*., 2015), and a 9 kb fragment containing the UAS-*Pex7* cassette at CII landing site attP40 (Bischof *et al*., 2007). Primers used:

MiMIC: 5’-TTGGAATCTGGAGCGTGGTGAG, 5’-ACTTGTACAGCTCGTCCATG
UAS-*Pex7*: 5’-CGGCTCTCGCAAATGCCAGCAG, 5’-CCTTCTAGACGACCAGATCAC

Primer specificity within the fly genome was verified using NCBI BLAST.

### Bioinformatic analysis

Percent identity matrices (PIMs) were generated by Clustal Omega multiple sequence alignment (Madeira *et al.,* 2019) using data acquired from the National Center for Biotechnology information (Geer *et al.,* 2010) and UniProt (The UniProt Consortium, 2018) databases. Protein sequences were aligned using the EMBOSS Needle application (Rice *et al*., 2000; McWilliam *et al*. 2013; Li *et al*., 2015). Protein domains were predicted using NCBI Conserved Domain Search (Marchler-Bauer *et al*., 2017).

### Data and custom reagent availability

All custom fly strains and plasmids noted herein are available upon request, subject to appropriate materials transfer agreements. The authors state all data necessary for confirming the conclusions presented are represented herein.

## Results

### Characterizing wildtype embryonic *Pex7* expression

A low-resolution assay of the relative level of endogenous *Pex* gene transcription in fixed *w^1118^* embryos was performed on embryos 2-4 and 4-8 h AEL (Figure 1). At 2-4 h, *Pex1* appeared most strongly in the yolk but by 4-8 h AEL had become ubiquitous (Figure 1A). The abundance of *Pex5* transcript appeared reduced at 4-8 h AEL while that of *Pex19* appeared to increase in the developing gut (Figure 1B). A similar, relatively less intense signal was also observed for the no-probe control. The pattern of cells with high levels of *Pex7* mRNA was different than other *Peroxins* at later time points.

**Figure 1.**
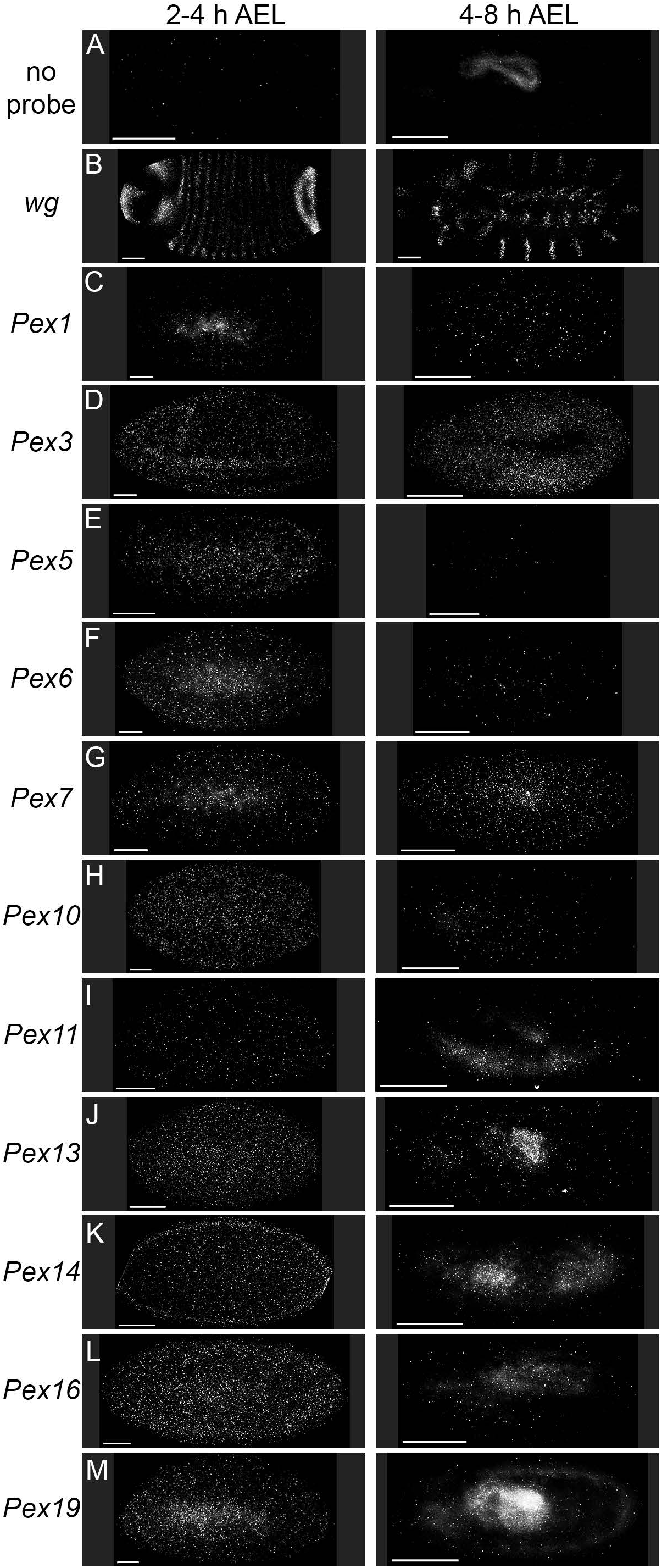
The transcription of *D. melanogaster Peroxin* genes during early embryogenesis as detected by FISH. (A) Exclusion of anti-sense probe allowed determination of tissue auto-fluorescence and served as the negative control. (B) The well-characterized mRNA localization pattern of *wg* (Baker, 1988) was used to validate our detection method and served as positive control. (C) *Pex1* appeared in the yolk after the MZT (left) then appeared ubiquitous, though faint, after cellularization (right). (D) *Pex3* mRNA appeared cortical during cephalization (left) and was strongly ubiquitous during gut formation (right). (E) *Pex5* mRNA appeared in the yolk and the cortex of the early syncytium (left) and by 4 h AEL became almost undetectable (right). (F) *Pex6* mRNA signal was very strong in the yolk and cortex of the syncytial blastoderm (left) and then became almost undetectable when gastrulation began (right). (G) *Pex7* mRNA signal localized with similar signal intensity and patterning to that of *Pex6* mRNA (left), and this signal retained its prevalence and pattern through the onset of gastrulation (right). (H) The syncytial pattern for *Pex10* mRNA localization was cortically ubiquitous (left) and did not appear in the yolk, then shifted to a broadly anterio-ventral pattern following cephalization (right). (I) *Pex11* mRNA appeared ubiquitous, like *Pex10* mRNA but lacking the same apparent number of signals (left). Following cephalization the pattern focused on the developing mesoderm and posterior midgut rudiment (right). (J) *Pex13* mRNA reported ubiquitously in the syncytial blastoderm with faint concentration visible in the yolk (left). During midgut development (right), *Pex13* was transcribed largely in the midgut with some cephalic and dorsal transcription as well. (K) During cellularization of the blastoderm, *Pex14* mRNA appeared ubiquitous within the yolk and localized to the outer cortical layer (left). Later, during gastrulation, *Pex14* mRNA appeared almost opposite in pattern to that of *Pex13* mRNA, localizing to the mesoderm without reporting from the posterior midgut rudiment or pole cells (right). (L) *Pex16* mRNA patterning was abundant and ubiquitous throughout following the MZT (left) and became restricted primarily to the elongating germ band and head primordium during midgut development (right). (M) After the MZT (left), *Pex19* mRNA localized to the yolk and appeared in cortical stripes reminiscent of a pair-rule gene (Nüsslein-Volhard and Wieschaus, 1980). This pattern altered after cephalization and became focused mainly on the posterior midgut rudiment and germ band (right). Scale bars = 50 µm, n = 3.

To more fully examine this pattern a complete spatio-temporal transcript localization profile of *Pex7* mRNA was performed on *w^1118^* embryos from 2-22 h AEL. Roughly 30 min after the maternal-zygotic transition (MZT) of gene transcription at ∼1.5 h AEL, foci of *Pex7* mRNA appeared in the yolk at the centre of the embryo (Figure 2A). These foci appeared cortically peri-nuclear just before syncytial cellularization (Figure 2B) then became ubiquitous following formation of the anterior-posterior axis and cephalization (Figure 2C). By 8 h AEL, *Pex7* transcription was restricted to the supraoesophageal ganglion and ventral nerve cord (Figure 2D; Hartenstein, 1993). The presumptive CNS localization pattern persisted, with increasing elaboration, until by 20 h AEL *Pex7* mRNA was found in subsets of cells on the CNS cortex (Figure 2E-G) that correlate spatially with persumptive neuroblasts (Hartenstein, 1993). Immediately prior to hatching at ∼22 h AEL, the pattern migrated into the optic lobe but appeared less elaborate overall (Figure 2H). The anterior/posterior fluorescence seen in the probe channel at ∼22 h AEL was non-specific fluorophore binding.

**Figure 2.**
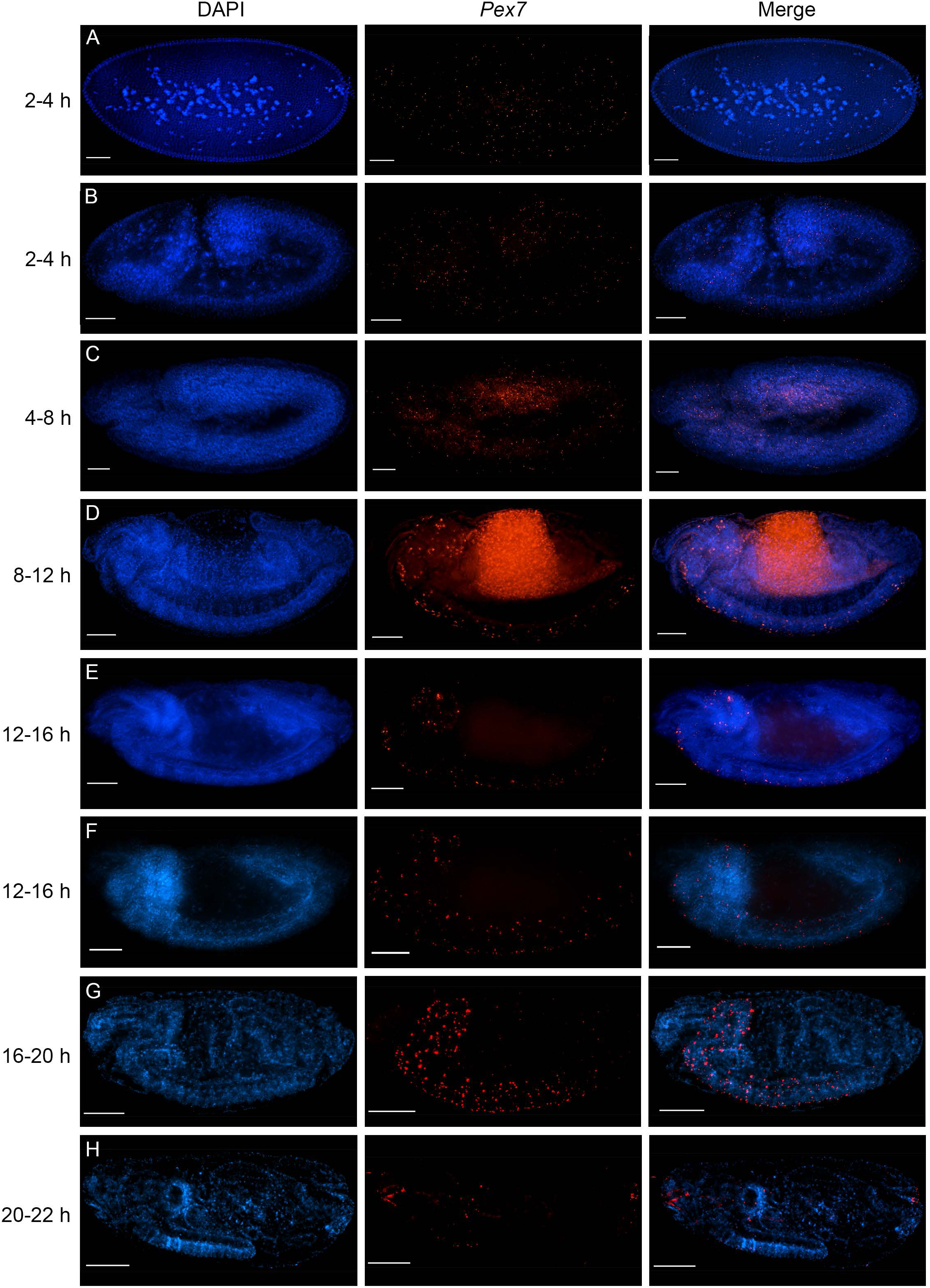
*Pex7* transcript localization during *D. melanogaster* embryogenesis. At 2-4 h AEL, localization was ubiquitous and did not overlap nuclei (A). Following formation of anterior-posterior axis and cephalization, transcription faded from the yolk (B). By 8 h AEL, *Pex7* transcription became restricted to the supraoesophageal ganglion and ventral nerve cord (E). The pattern was maintained until hatching (F-H) though some diffusion was observed by ∼20 h AEL (H). Gut auto-fluorescence in C-F was an artifact of primary antibody detection. Scale bars = 50 µm, n = 3.

As *Pex7* expression includes translation of a cytosolic polypeptide in species that have the gene, we raised an antibody in rabbits targeting residues 46-65 of the predicted *D. melanogaster* Pex7 protein. These residues occur immediately upstream of the first predicted WD40 domain repeat in a soluble β-sheet motif (Figure 9C). A GST-Pex7 chimera was grown in *E. coli* strain BL-21 and lysate was blotted with αGST and αPex7. The resultant bands overlapped at ∼66 kDa (data not shown), and as the GST tag blots at 28 kDa (ThermoFisher) the apparent mass of Pex7 was deduced to be ∼38 kDa, close to its predicted mass of 37 kDa. A 3xFLAG-*Pex7* chimera was then grown in Schneider 2 (S2) cells and lysates were blotted with αFLAG and αPex7. The resultant bands overlapped at ∼48 kDa (Figure 3A-C). αPex7 was then blotted along with βTub56D antibody E9 on lysates of *w^1118^* embryos (Figure 3D-H). Endogenous Pex7 appeared at ∼40 kDa (lower bands) and E9 reported at 56 kDa (upper bands) in all blots, without overlap. The apparent ∼2 kDa bandshift between chimeric and endogenous Pex7 was attributed to post-translational modification of the endogenous protein.

**Figure 3.**
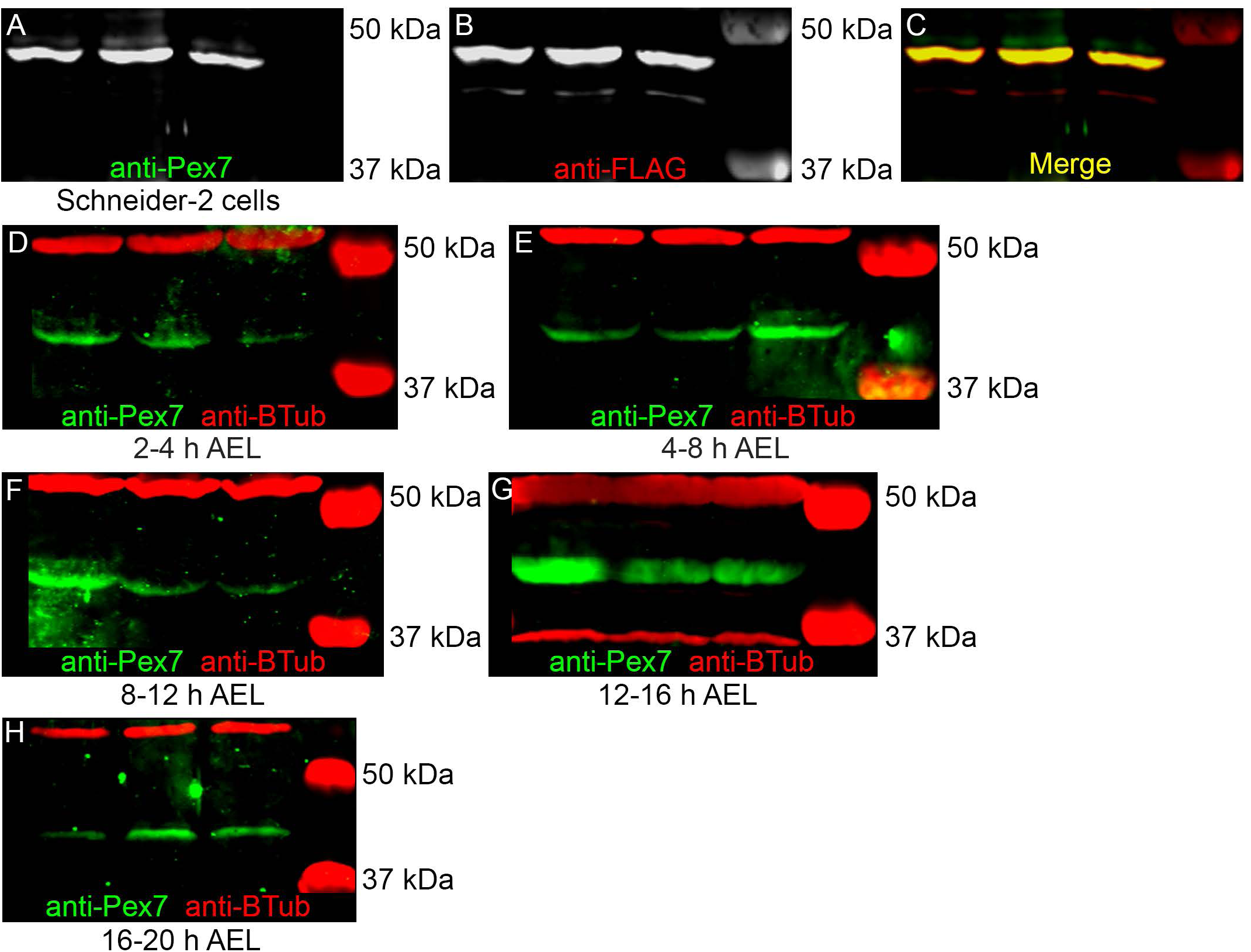
Translation of *D. melanogaster Pex7*. (A-C) Chimeric 3xFLAG-Pex7 expressed in *D. melanogaster* Schneider-2 (S2) cells revealed overlap of αPex7 and αFLAG reporters at ∼48 kDa. D-H: Lysates of *w^1118^* embryos collected at 2-4 h (D), 4-8 h (E), 8-12 h (F), 12-16 h (G) and 16-20 h (H) AEL were blotted with αPex7 and α-βTubulin, which reported at the apparent sizes of ∼40 kDa and ∼56 kDa, respectively, without overlap (n=3).

Beginning at 2 h AEL, Pex7 signal appears in all cells (Figure 4A). At the onset of gastrulation around 190 minutes after egg laying (min AEL), Pex7 IF signal was below detectable levels (Figure 4B) until germ band elongation completed (220 min AEL) when Pex7 again appeared ubiquitous. This pattern maintained through slow germ band elongation, gnathal/clypeolabral lobe formation and germ band retraction at 440 min AEL (Figure 4C; Hartenstein, 1993). During this period Pex7 did not co-localize with Wingless (Wg), a paracrine/autocrine ligand that contributes to segment polarity and the morphogenesis of neuromuscular junctions (Nüsslein-Volhard and Wieschaus, 1980; Patel, Schafer *et al*., 1989; Volk and Vijay Raghavan, 1994; Klingensmith and Nusse, 1994). From 440-580 min AEL, when the optic lobes invaginate and the nervous system differentiates from progenitor neuroblasts, Pex7 localized para-segmentally to cells adjacent to those marked by the pan-neural antibody marker 9F8A9, which targets Embryonic Lethal Abnormal Vision (αElav), again without overlapping (Figure 4D). By 12 h AEL the pattern had elaborated para-segmentally into five small distinct branches (Figure 4E) and two foci adjacent to the ventral nerve cord (Figure 4E, middle panel, white arrowhead). This pattern did not overlap with the CNS/PNS marker antibody BP 104, which targets Neuroglian (αNrg), or the glial antibody marker 8D12, which targets Reversed Polarity (αRepo, Figure 4F). By 16 h AEL the αPex7 pattern appeared similar to that of its transcript, localizing to a subset of cells in the central nervous system with some signal in nearby cells and the cuticle (Figure 4G). By 20 h the αPex7 pattern diffused though concentration in the cuticle and adjacent to the CNS was observed (Figure 4H).

**Figure 4.**
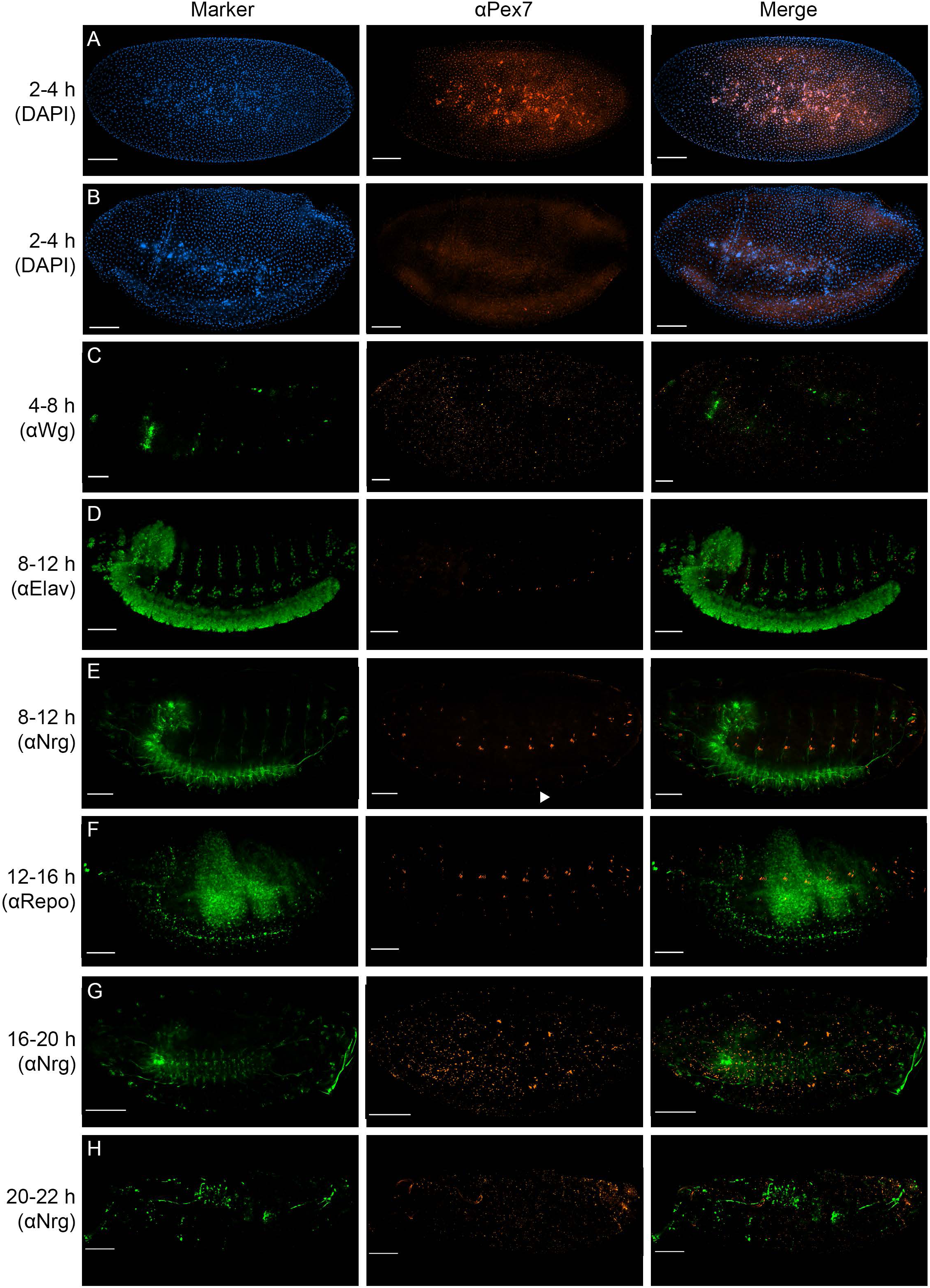
Embryonic Pex7 translation. Pex7 can be detected in a restricted subset of embryonic cells by immunoflourecence (IF) detection. The Pex7 IF signal overlapped that of nuclei in the embryo core (A) which was maintained until gastrulation at 3 h AEL when it approached the detection threshold (B). The report then became ubiquitous when rapid germ band elongation began and persisted until 7.5 h AEL without overlapping the neuromuscular junction marker αWg (D). At 8 h AEL Pex7 was restricted para-segmentally to cells adjacent to differentiating neurons, marked by αElav (E). By 12 h AEL the pattern elaborated into distinct bands and migrated ventrally to the nerve cord without overlapping the nervous system marker αNrg (F, middle panel, white arrowhead). Expansion continued dorso-ventrally within each para-segment through 16 h AEL, without overlapping the glial reporter αRepo (G). Pex7 then diffused at ∼20 h AEL with concentrations at the CNS and cuticle (H). Gut auto-fluorescence in G and extreme dorso-ventral autofluorescence in H were detection artifacts. Scale bars = 50 µm, n = 3.

Our qualitative *Pex7* data suggested its embryonic expression was regulated temporally and therefore either or both of its transcript and polypeptide varied in abundance depending on developmental stage. To investigate this, relative embryonic transcript and protein abundance were assayed via qPCR and fluorescence-based western blotting, respectively (Figure 5).

**Figure 5.**
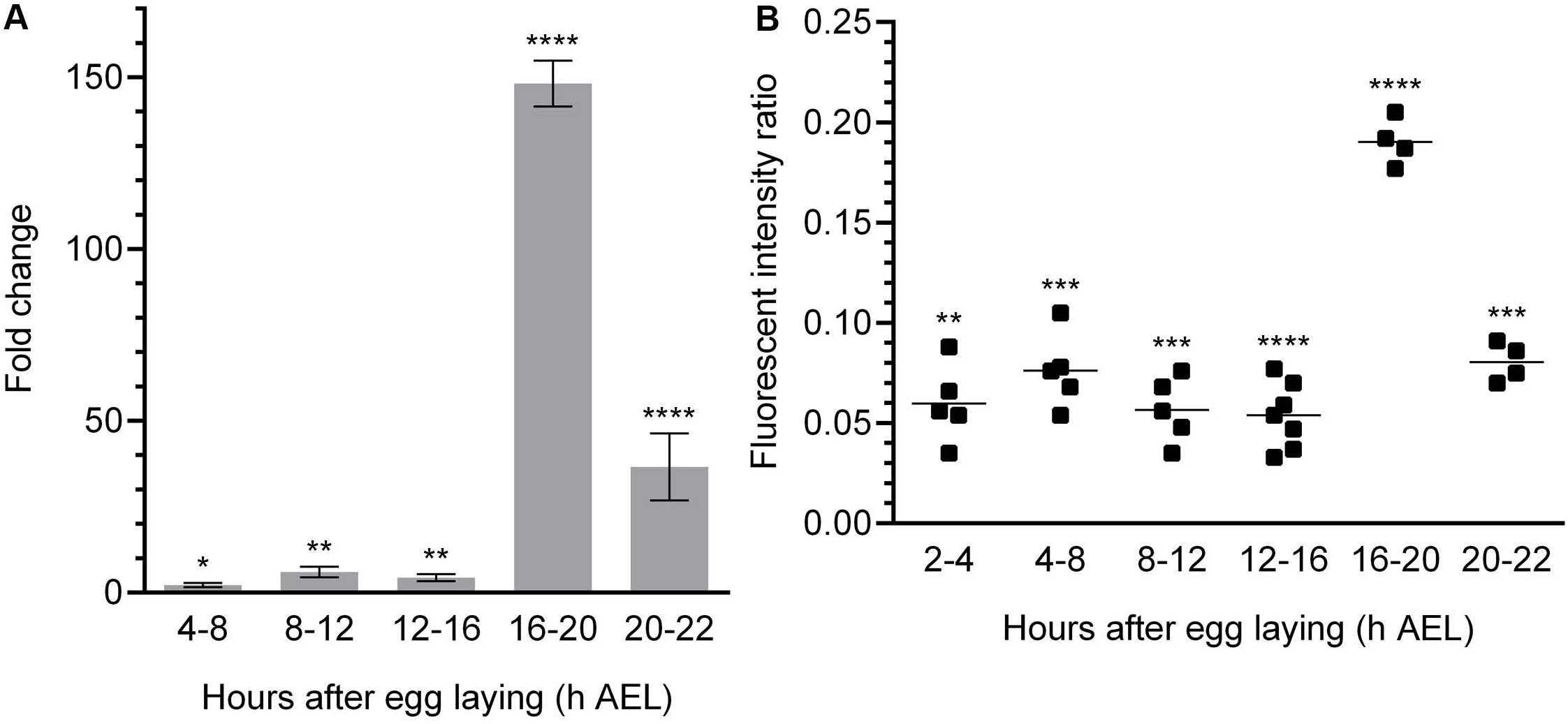
Quantitation of wild-type embryonic *Pex7* expression. Using *w^1118^* embryos, *Pex7* transcript abundance was quantified relative to *Act5C* and Pex7 polypeptide abundance was quantified relative to βTubulin56D. (A) qRT-PCR of *Pex7* transcript abundance was measured relative to *Act5C* in embryos collected 2-4 h AEL, then used to compute relative fold-change in abundance at various time points spanning embryogenesis (n = 3, two primer pairs each). Data presented are mean fold-change (n = 4 comparisons) ± SD. *P* values were determined by comparing ΔCt values at each time point to the 2-4 h AEL baseline (Table 1). (B) Relative Pex7 polypeptide abundance was measured by comparing the same-lane Pex7:βTubulin fluorescent intensity ratios in western blots of staged embryo lysates (n≥4). *P* values were determined by comparing pooled replicate values at each time point to a null hypothesis of 0. * *P*≤0.02, ** *P≤*0.01, *** *P≤*0.001, **** *P≤*0.0001.

**Table 1.** Summary of qRT-PCR data

| Mean C<sub>t</sub> Scores | | | Mean $\Delta C_t$: *Pex7* vs *Act5C* | | | | |
| --- | --- | --- | --- | --- | --- | --- | --- |
| Name | Mean C<sub>t</sub> | St Dev | Pair | $\Delta C_t$ | St Dev | $\Delta\Delta C_t$ | $2^{-\Delta\Delta C_t}$ |
| 2-4 h AEL |  |  |  |  |  |  |  |
| *Pex7-1* | 36.49 | 0.12 | *Pex7-1/Act5C-2* | 6.45 | 0.03 | - | - |
| *Pex7-3* | 35.76 | 0.19 | *Pex7-1/Act5C-3* | 5.90 | 0.16 | - | - |
| *Act5C-2* | 30.04 | 0.18 | *Pex7-3/Act5C-2* | 5.72 | 0.01 | - | - |
| *Act5C-3* | 30.59 | 0.43 | *Pex7-3/Act5C-3* | 5.17 | 0.12 | - | - |
| 4-8 h AEL |  |  |  |  |  |  |  |
| *Pex7-1* | 34.02 | 0.34 | *Pex7-1/Act5C-2* | 4.87 | 0.13 | -1.58 | 2.99 |
| *Pex7-3* | 33.54 | 0.11 | *Pex7-1/Act5C-3* | 5.10 | 0.06 | -0.80 | 1.74 |
| *Act5C-2* | 29.15 | 0.08 | *Pex7-3/Act5C-2* | 4.39 | 0.02 | -1.33 | 2.51 |
| *Act5C-3* | 28.92 | 0.46 | *Pex7-3/Act5C-3* | 4.62 | 0.18 | -0.55 | 1.46 |
| 8-12 h AEL |  |  |  |  |  |  |  |
| *Pex7-1* | 35.87 | 0.36 | *Pex7-1/Act5C-2* | 4.42 | 0.04 | -2.03 | 4.08 |
| *Pex7-3* | 34.48 | 0.09 | *Pex7-1/Act5C-3* | 3.52 | 0.03 | -2.38 | 5.21 |
| *Act5C-2* | 31.45 | 0.29 | *Pex7-3/Act5C-2* | 3.03 | 0.10 | -2.69 | 6.45 |
| *Act5C-3* | 32.35 | 0.42 | *Pex7-3/Act5C-3* | 2.13 | 0.16 | -3.04 | 8.22 |
| 12-16 h AEL |  |  |  |  |  |  |  |
| *Pex7-1* | 36.35 | 0.22 | *Pex7-1/Act5C-2* | 4.82 | 0.09 | -1.63 | 3.10 |
| *Pex7-3* | 35.34 | 0.21 | *Pex7-1/Act5C-3* | 3.64 | 0.12 | -2.26 | 4.79 |
| *Act5C-2* | 31.53 | 0.05 | *Pex7-3/Act5C-2* | 3.81 | 0.08 | -1.91 | 3.76 |
| *Act5C-3* | 32.71 | 0.46 | *Pex7-3/Act5C-3* | 2.63 | 0.13 | -2.54 | 5.82 |
| 16-20 h AEL |  |  |  |  |  |  |  |
| *Pex7-1* | 33.08 | 0.45 | *Pex7-1/Act5C-2* | -0.67 | 0.12 | -7.12 | 139.10 |
| *Pex7-3* | 32.28 | 0.21 | *Pex7-1/Act5C-3* | -1.33 | 0.13 | -7.23 | 150.12 |
| *Act5C-2* | 33.75 | 0.68 | *Pex7-3/Act5C-2* | -1.47 | 0.24 | -7.19 | 146.02 |
| *Act5C-3* | 34.41 | 0.19 | *Pex7-3/Act5C-3* | -2.13 | 0.01 | -7.30 | 157.59 |
| 20-22 h AEL |  |  |  |  |  |  |  |
| *Pex7-1* | 32.22 | 0.54 | *Pex7-1/Act5C-2* | 0.90 | 0.06 | -5.55 | 46.85 |
| *Pex7-3* | 31.52 | 0.18 | *Pex7-1/Act5C-3* | 1.14 | 0.13 | -4.76 | 27.10 |
| *Act5C-2* | 31.32 | 0.65 | *Pex7-3/Act5C-2* | 0.20 | 0.24 | -5.52 | 45.89 |
| *Act5C-3* | 31.08 | 0.29 | *Pex7-3/Act5C-3* | 0.44 | 0.06 | -4.73 | 26.54 |

Table 1 continued
|  | Mean | St Dev | P value |
| --- | --- | --- | --- |
| | $2^{-\Delta\Delta C_t}$ | $2^{-\Delta\Delta C_t}$ | $\Delta C_t$ |
| 4-8 h AEL | 2.18 | 0.607 | 1.855E-02 |
| 8-12 h AEL | 5.99 | 1.538 | 6.685E-03 |
| 12-16 h AEL | 4.36 | 1.033 | 1.087E-02 |
| 16-20 h AEL | 148.21 | 6.696 | 2.147E-06 |
| 20-22 h AEL | 36.59 | 9.784 | 7.482E-06 |

Relative *Pex7* mRNA abundance in *w^1118^* embryos was measured relative to that of *Act5C* using the ΔΔC_t_ method as described previously (Livak and Schmittgen, 2001) and presented herein as mean ± SD. Two primer pairs for each gene passed verification (Table 1), from which the total mean fold change and standard deviation were calculated and plotted. The initial ΔC_t_ of 5.17 ± 0.12 at 2-4 h (Table 1) served as baseline and indicated *Pex7* was transcribed at this stage. Mean fold-change in transcript abundance increased by 2.18 ± 0.61 at 4-8 h AEL (*P* = 0.0186) and 5.99 ± 1.54 by 8-12 h AEL (*P* = 0.00669), indicating a steady increase in transcript abundance as it localized to the CNS (Figure 2D). At 12-16 h the fold-change still increased but slowed to 4.36 ± 1.03 (*P* = 0.0109), a downward trend observed for all primer pairs (Table 1). At 16-20 h AEL the fold-change plateaued at 148.21 ± 6.70 (*P* = 2.15 x 10^−6^), when the most elaborate *Pex7* mRNA CNS patterning was observed (Figure 2G), then dropped to 36.59 ± 9.78 by 20-22 h AEL (*P* = 7.48 x 10^−6^). Variability in fold change between primers used was attributed to primer efficiency as the trend in fold-change did not differ (Table 1; Livak and Schmittgen, 2001).

A difference was observed in the localization of *Pex7* transcript and protein before 16 h AEL, suggesting differential regulation. To test this, the abundance of Pex7 protein was measured relative to βTub56D in western blots of embryo lysates (Figure 5B, Table 2), normalized to replicate-specific background fluorescence, and presented herein as mean ± SD. At 2-4 h AEL the ratio was a low but detectable 0.060 ± 0.017 (*P* = 0.022). The mean ratio increased to 0.076 ± 0.017 at 4-8 h AEL (*P* = 0.00084), trending in agreement with its transcript, then dropped to 0.057 ± 0.014 at 8-12 h AEL (*P* = 0.0014) and further to 0.054 ± 0.015 at 12-16 h AEL (P = 0.00013). This contrasted with the qRT-PCR fold-change data, which was above the 2-4 h AEL baseline at these stages (Figure 5A). The mean ratio rose sharply at 16 h AEL to 0.190 ± 0.010 (*P* = 6.6 x 10^−5^) then dropped to 0.081 ± 0.008 by 20 h AEL (P = 4.4 x 10^−4^). The temporal pattern of relative Pex7 abundance supported our qualitative expression data.

**Table 2.**
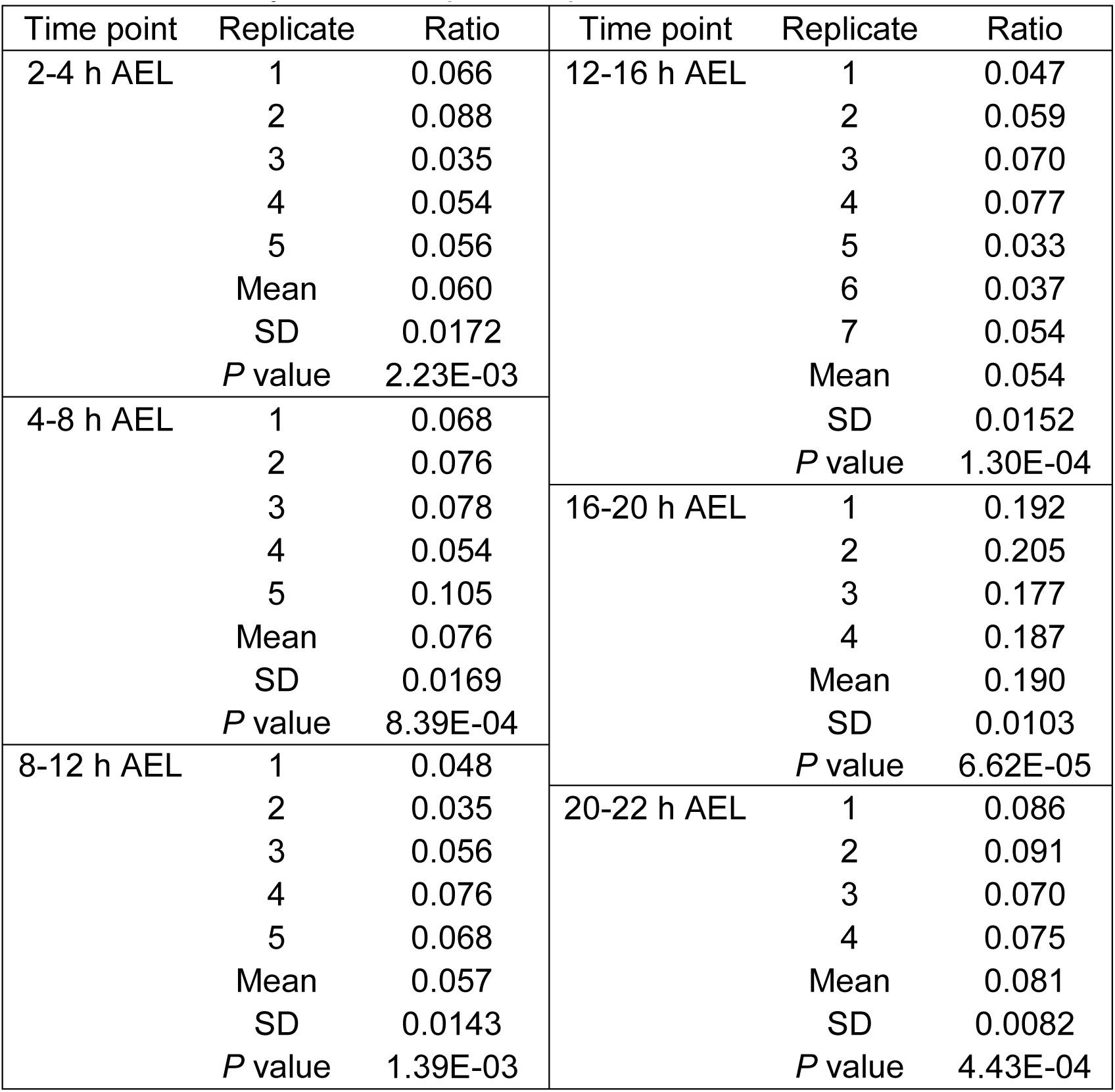
Summary of relative protein quantification

**Table 3.**
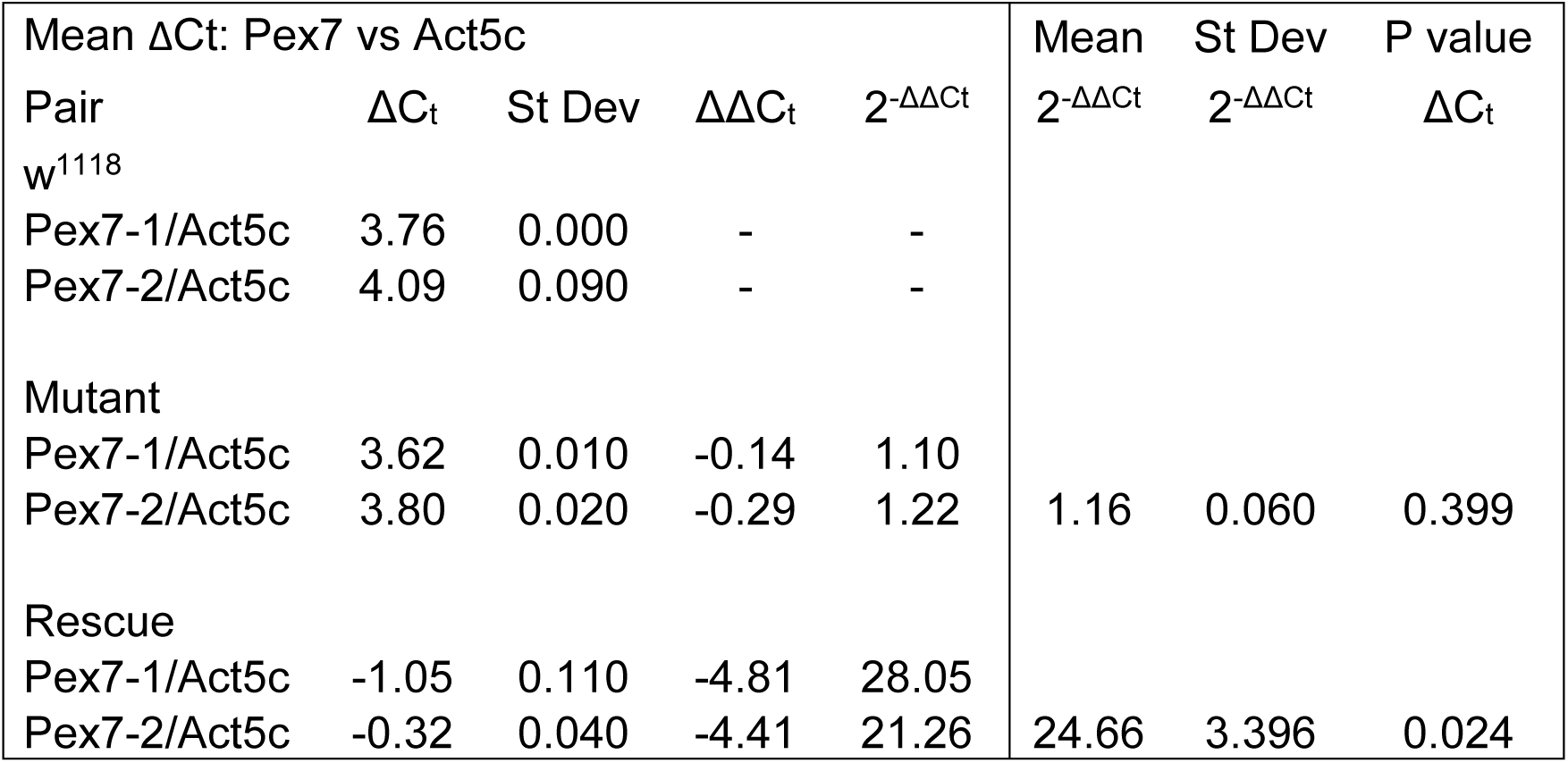
Summary of adult qRT-PCR data

### Forward *Pex7* mutagenesis screen

The dynamic change in the cells transcribing *Pex7* and the similar changes in relative protein expression during embryonic development suggested Pex7 had a functional role. Recently, homozygous adult *Pex7^MiMIC^* mutants were reported to have difficulty with negative geotaxis and displayed a sub-lethal, semi-pharate phenotype (Di Cara *et al*., 2019; Linderman *et al.,* 2012). We hypothesized these phenotypes arose due to affected nervous system development during embryogenesis. A targeted mutation adult survivorship assay was performed using the Gal4-UAS system paired with CRISPR-mediated gene knockout (TRiP-CRISPR KO, TRiP-KO). Briefly, parent strains bearing complementary TRiP-KO genomic cassettes were mated to produce progeny in which somatic *Pex7* knockout was directed to all developing neuroblasts (*elav*-Gal4), and the developing peripheral nervous system (*pnr*-Gal4) (Gratz *et al.,* 2015; Lin *et al.,* 2015). Somatic *Pex14* knockout was used as a positive control as this would cause loss of Pex5-mediated peroxisome enzyme import (Faust *et al*., 2012; Fukiji *et al*., 2014; Baron *et al*., 2016). The negative control strain (Scramble) targeted the *N. crassa* gene QUAS which has no fly homolog (Gratz *et al.,* 2015). Mutant adult fly counts were lower than control for *elav>Pex7* and *elav>14* TRiP-KO, as were total adults, while neither *pnr*>*Pex7* nor *pnr>Pex14* knockout were effective (Figure 6A-B). Statistical phenotypic significance was assessed by comparing the mutant:total ratio of surviving adults. Mutant adults were significantly reduced for *elav>Pex7* alone (*P* = 0.055, Figure 6C). Male:total adult ratios did not vary significantly indicating no sex-specific effect occurred (Figure 6D). As adults continued to eclose from these crosses, this confirmed the novel tissue-specific *Pex7* knockout phenotype was semi-lethal. To isolate the specific stage at which loss of Pex7 began to affect survival, total progeny eggs laid and hatched from TRiP-KO crosses were scored. *elav*>*Pex7* KO resulted in significantly fewer larva 4 d after egg laying (*P* = 0.047, Figure 6E) and a reduced larva:egg ratio (*P* = 0.029, Figure 6F), indicating the semi-lethal *Pex7* phenotype onset was during embryogenesis.

**Figure 6.**
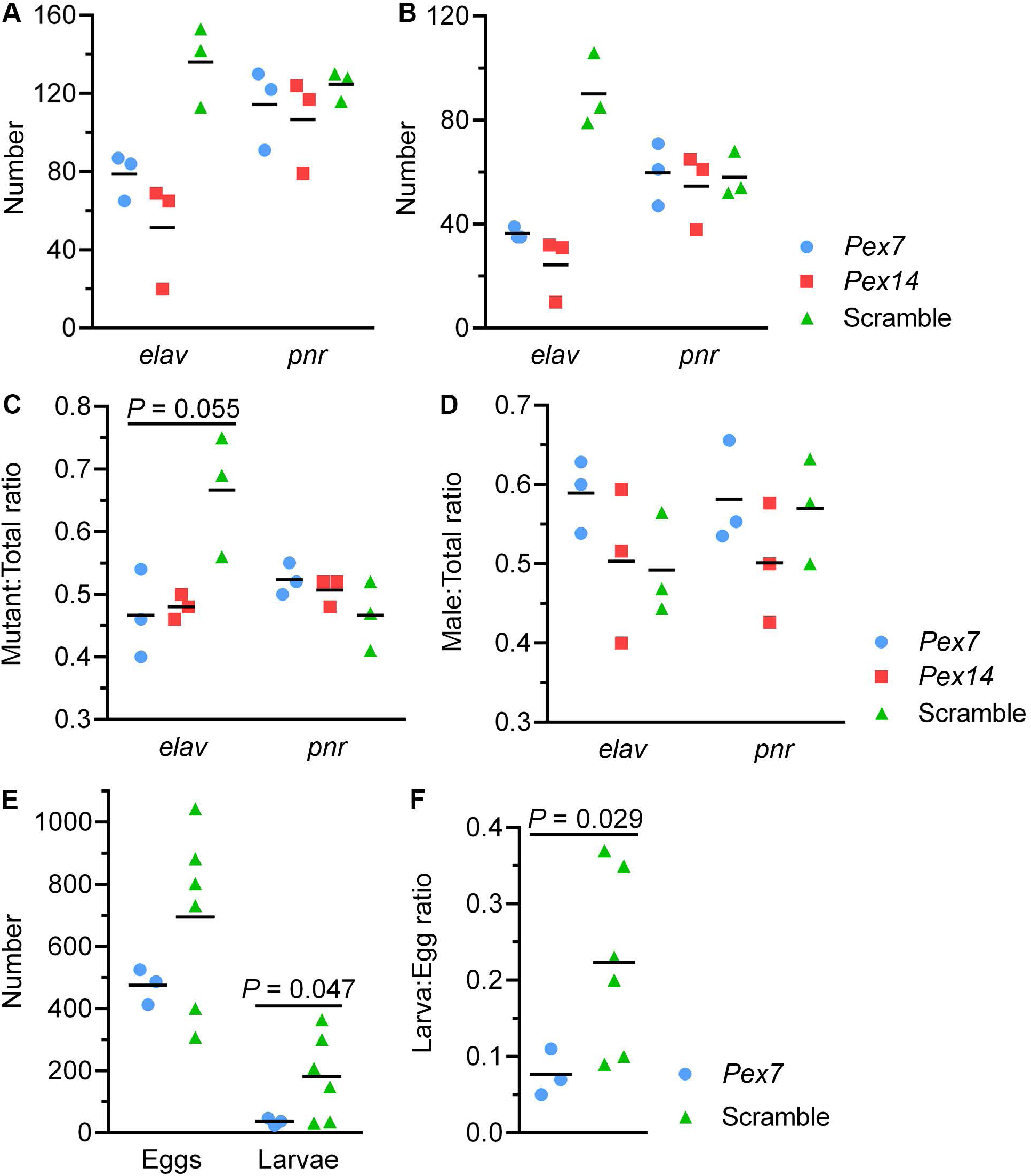
Somatic knockout of *Pex7* in differentiating neuroblasts impaired hatching. (A) *elav*>*Pex7* and *elav*>*Pex14* TRiP-KO counted significantly reduced mean adult mutants whereas *pnr*>*Pex7* and *pnr*>*Pex14* did not. (B-C) Adult mutant sex ratios did not differ significantly for either *elav*-driven (B) or *pnr*-driven (C) *Pex7* knockout. D-F: Egg tallies (D), larva tallies (E) and larva:egg ratios (F) of somatic *elav*>*Pex7* TRiP-KO mutants. Somatic *elav*>*Pex7* mutation significantly reduced larvae counts (E, *P* = 0.055) with concomitant reduction in larva:egg ratio (F, *P* = 0.029).

To examine the effect of over-expressing *Pex7*, ubiquitous and neuroblast-specific targeting was mediated by Gal4-*Act5C* and Gal4-*elav* cassettes, respectively. Somatic over-expression of *Pex1* was used as a Peroxin-positive control (Mast *et al.,* 2011) and progeny bearing un-driven UAS-*Pex7* was the negative control. Counts of total adult and total mutant progeny were reduced for *elav>Pex7* and *Act5C>Pex7* over-expression (Figure 7A and B, respectively) with the presence of adult mutants indicating an effect on viability. The mutant:total adult ratio was significantly reduced for *elav*>*Pex7* (*P* = 0.053), *Act5C*>*Pex7* (*P* = 0.026) and *Act5C*>*Pex1* (*P* = 0.0017) over-expression (Figure 7C), with the phenotype being significantly more pronounced for *Pex1* than *Pex7* under *Act5C* (*P* = 0.015). *Pex7* over-expression had a milder effect on total adult counts and total mutant adult counts than *Pex1* however the distribution of *elav>Pex7* replicates was bi-modal and its mean progeny count was lower. Male:total adult ratios were unaffected indicating no sex-specific effect was present (Figure 7D). The *elav>Pex7* replicates with the lowest adult counts also had male mutant progeny outnumbering female progeny by 2:1 (Figure 7D) while the sex ratios of non-mutant progeny of the same cross were not affected. Egg and larva counts revealed a reduction in eggs laid for *Pex7* under both drivers, in turn impacting total larvae hatched, without also affecting the larva:egg ratio (Figure 7E-F). Of note, parent females lacked the cassettes necessary to manifest somatic over-expression, un-driven UAS-*Pex7* did not replicate the phenotype and, when employed as part of TRiP-KO, the mere presence of Gal4-UAS cassettes did not significantly affect egg or adult progeny counts (Figure 6).

**Figure 7.**
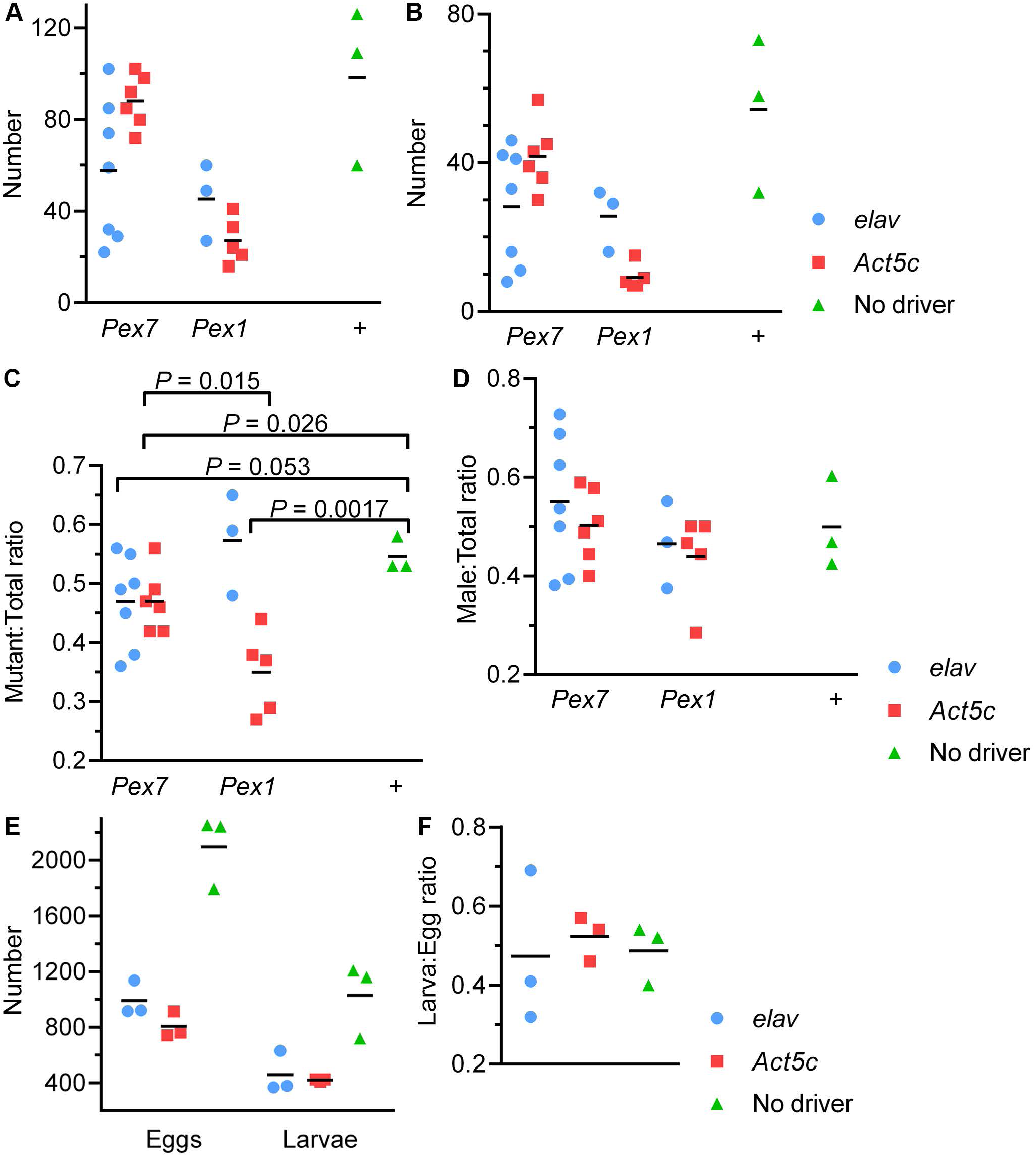
Somatic *Pex7* over-expression affected viability without impairing hatching. Promoters of either *elav* or *Act5C* were used to direct over-expression (OE) of Pex7 via Gal4-UAS. (A-B) Mean total adults and mean total mutant adults were reduced by both drivers. (C-D) *Act5C>Pex7* and *elav>Pex7* had significantly reduced mutant:total adult ratios (*P* = 0.053 and 0.026, respectively). This phenotype was also observed for *Act5C>Pex1* (*P* = 0.0017) and was significantly stronger than for *Act5C>Pex7* (*P* = 0.015). No effect on adult sex ratio was observed. (E-F) Total eggs laid were reduced, also reducing larva hatched, without affecting larva:egg ratio.

Finally, to determine if the neuronal-lineage specific phenotypes associated with the *Pex7^MiMIC^* were caused by loss of Pex7 in these tissues, a fly strain that expressed *Pex7* in the neuroblasts of homozygous *Pex7* mutants (*elav*>*Pex7*; *Pex7^MiMIC^*/*Pex7^MiMIC^*) was examined. No significant difference was observed in total adult progeny (Figure 8A), sex ratios (Figure 8B) or total eggs laid and larvae hatched (Figure 8C) between rescued and un-rescued mutants, and the larva:egg ratio was significantly reduced under the rescue condition (*P* = 0.037, Figure 8D), a sub-lethal phenotype similar to that observed under somatic knockout. The genomic MiMIC insert and UAS-*Pex7* cassette were verified in rescued adult mutant gDNA by PCR, and *Pex7* transcription levels were verified in rescued and un-rescued adults by qRT-PCR (Figure 8E-G), presented herein as mean ± SD. *Pex7* transcript abundance in mutants was roughly equal to that of control w^1118^ adults, with a mean fold-change difference of 1.16 ± 0.06, and that of rescued mutants was 24.66 ± 3.40 (*P* = 0.024, Figure 8G).

**Figure 8.**
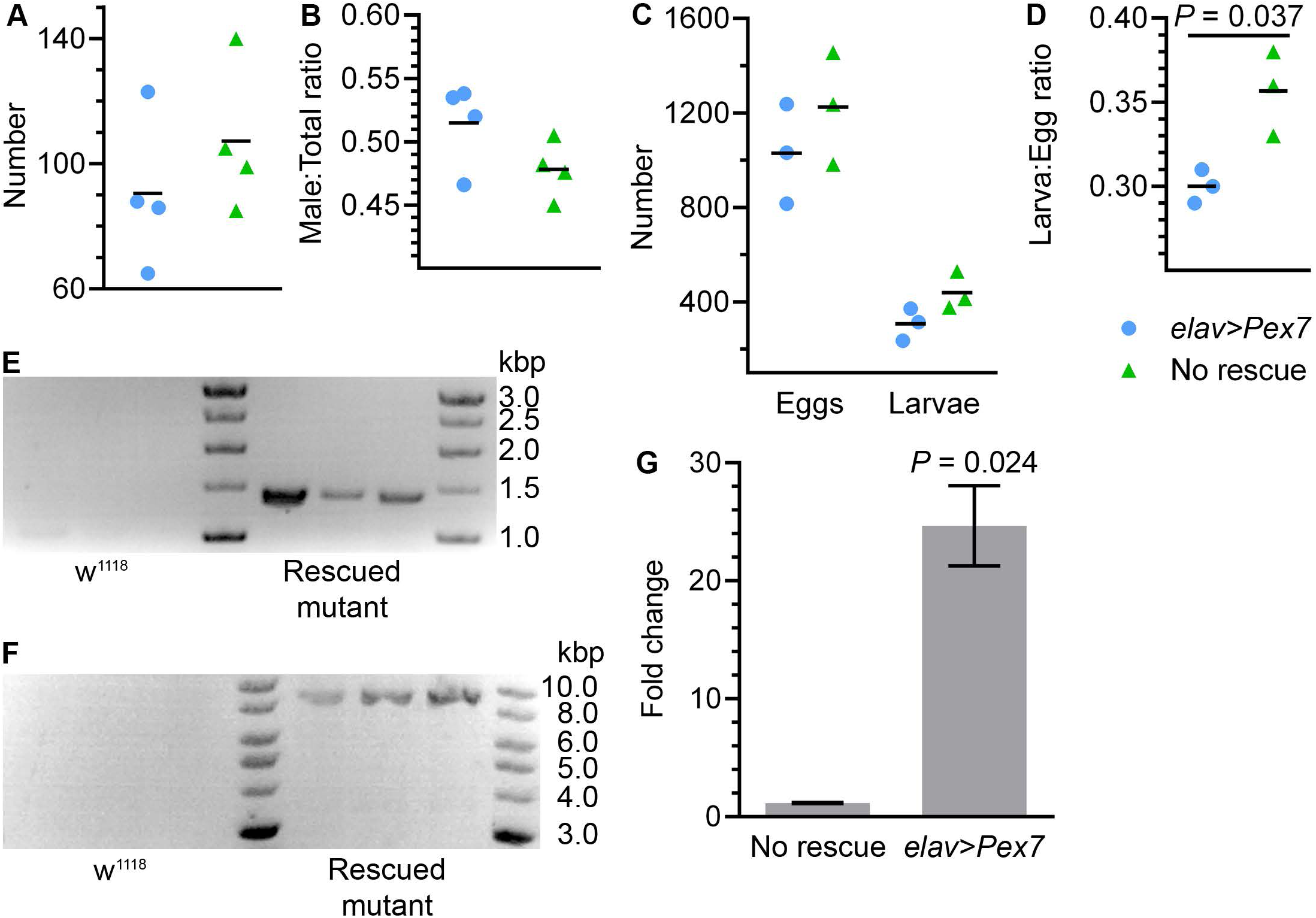
Targeted *Pex7* rescue in homozygous *Pex7* mutants impaired hatching. (A-B) Neither total counts nor adult sex ratios differed significantly between rescued and non-rescued adults. (C-D) Total eggs laid and larvae hatched did not differ significantly between rescued and non-rescued adults, however the rescued mutants had a significantly lower hatching rate (*P* = 0.037). (E-F) The presence of the MiMIC insert (C) and UAS-*Pex7* cassette (F) were verified by PCR of rescued adult gDNA. (G) *Pex7* transcript abundance was assayed in rescued and non-rescued adults by qRT-PCR and reported as fold-change difference from strain *w*^1118^ (n = 3). Rescued mutants had significantly increased *Pex7* expression (*P* = 0.024).

## Discussion and Conclusions

To date, screens of fly *Peroxins* have focused mainly on larval and adult phenotypes. For example, third instar *pex3* larvae were smaller and contained less ceramide than their rescued counterparts, and *Dicer*-enhanced *Mef2>Pex3* RNAi resulted in failed/difficult eclosure (semi-pharate) and reduced adult survivorship (Faust et al., 2014). *pex5* adults displayed similar phenotypes, as well as highly disorganized neural, glial and muscular patterns during embryogenesis and altered larval very long-chain fatty acid (VLCA) species abundance (Di Cara *et al.,* 2019). *pex7* larvae had reduced CNS size and increased neuro-specific apoptosis activity that was rescued somewhat by *nos>Pex7*. Neuro-specific apoptosis was also elevated in *pex7* embryos (Di Cara *et al.,* 2019). These findings strongly suggested global *Peroxin* mutation resulted in tissue-specifc effects, which we explored in the present study. During blastoderm embryo development *Peroxin* mRNAs are distributed relatively equally in the syncytial embryo and following cellularization (Figure 1). During early gastrulation, differential *Peroxin* mRNA localization was observed. *Pex1* and *Pex6* mRNA reported similar patterns at 4-8 h AEL, inferring conservation of a relationship their homologs have in yeast and human (Motley *et al.,* 2015; Ciniawsky S *et al.,* 2015; Tamura *et al.,* 2006). *Pex3* mRNA was ubiquitous regardless of stage while that of *Pex5* was almost undetectable during gastrulation, suggesting most of the Pex5 required during this milestone was translated beforehand. *Pex10* mRNA had a similar pattern to that of *Pex5*, suggesting their transcription was co-ordinated and inferring conservation of their relationship (Chang *et al.,* 1999). The restriction of *Pex11* mRNA to the germ band and posterior midgut rudiment at 4-8 h AEL, differing from the strong midgut signal observed for *Pex19* mRNA in the same period, suggests *Pex11* is involved somehow in developing tissue with high lipid metabolism needs such as the nervous system and the gut, but not predominantly in the gut like *Pex19*. The overall patterning of *Pex19* mRNA at 4-8 h AEL appeared similar to that of *Pex13* mRNA, which is interesting given their conserved functions are not part of the same process; *Pex19* is a chaperone that regulates the import of peroxisome membrane proteins into peroxisomes during early peroxisome biogenesis (Sacksteder *et al.,* 2000) and *Pex13* forms part of the PTS1 import translocon that facilitates import of cargo-bearing Pex5 (Dias *et al.,* 2017).

Interestingly, *Pex19* mRNA patterning did not match that of *Pex3* mRNA, which suggests their relationship is not conserved (Götte *et al.,* 1998). Our screen did not identify embryonic tissue specificity for *Pex3* mRNA.

Variation in human PEX7, either homozygous or compound heterozygous, manifests as RCDP type 1 (RCDP1) which leads to defects during embryo development (Braverman *et al*., 2012; Krakow *et al*., 2003). The qualitative (Figures 2 and 4) and quantitative (Figure 5, Tables 1 and 2) analyses of *Pex7* in strain *w^1118^* embryos affirmed that Pex7 is transcribed/translated at relatively high levels in highly restricted subset of developing cells, likely of neural linage. Building on this, we observed that reducing Pex7 levels only in these tissues (*elav>Pex7* TRiP-KO) affected survival to adulthood (Figure 6). This effect was also observed to a lesser degree for *elav>Pex14* TRiP-KO. This suggests that there is a general requirement for peroxisomes in these cell linages, but that *Pex7* is needed specifically by neuroblasts as its mutant phenotype was stronger in those cells. The relative increase in overall *Pex7* expression, as well as increased expression in presumptive neuronal linages at 16-20 h AEL (Figure 5) are ascribed to the developmental needs of neuroblasts on the CNS cortex, given that *elav-*targeted *Pex7* CRISPR induced somatic mutation reduces viability by impairing hatching ability and impairing escaper co-ordination (Di Cara *et al*., 2019). Further, the similarity in phenotype nature, magnitude and specificity between *Pex7* and *Pex14* TRiP-KO, and *Pex1* and *Pex7* over-expression, suggest *Drosophila Pex7* is involved in peroxisome-related processes as it is in yeast and mammals (Neuhaus *et al*., 2014; Mast *et al.,* 2011; Albertini *et al*., 1997; Braverman *et al*., 1997; Reuber *et al*., 1997; Erdmann *et al*., 1991).

The negative data from *pnr>Pex7* and *pnr>Pex14* TRiP-KO (Figure 6), supported by our *in situ* data (Figure 1), shows that this requirement for *Pex7* is likely specific to the central nervous system. The acute requirement for Pex7 in CNS was also supported by our TRiP-KO data, which identified the sensitivity of the developing CNS to *Pex7* loss. We observed *elav>Pex7* over-expression reduced mean total adult and mean mutant adult progeny more severely than *Act5C>Pex7*, while the effect was reversed for *Pex1* over-expression (Figure 7). This supports a CNS-specific function for Pex7 and suggested it had an ideal range of expression depending on developmental stage. The strong reduction in total eggs laid for both gene mutations was surprising (Figure 7C) given the P generation was not known to have any maternal effect or spermatogenesis mutations reducing their fertility and that the assay specifically induced somatic mutation in progeny only. A similar but non-significant effect on egg laying was observed in our somatic TRiP-KO assay (Figure 6E), however the control data included counts within the same range while maintaining a higher larva:egg ratio (Figure 6F). This indicates, in crosses bearing a higher mutation load, that the observation is not likely due to a P-generation fertility effect. Moreover, as the over-expression control progeny bore un-driven UAS-*Pex7* and a similar phenotype was not observed for counts of *pnr>Pex7* TRiP-KO progeny (Figure 6A-D), we discounted cryptic Gal4-UAS fertility variables and posited the reduced egg counts were due to non-specific effects of somatic *Peroxin* over-expression, gene-specific effects notwithstanding. A mechanism for how directed somatic *Peroxin* over-expression affects apparent parental fertility needs to be examined further, considering zygotic gene expression begins ∼2 h after the egg is laid (Anderson and Lengyel, 1979). The sub-lethal phenotype we observed following *elav> Pex7* rescue was similar to that of *elav>Pex7* TRiP-KO, suggesting an ideal range of *Pex7* expression exists in developing neuroblasts.

### Functional conservation

In human, *in utero*/neonatal clinical diagnosis of RCDP1 relies on identification of skeletal defects, inappropriate or missing ossification of cartilage, and biochemical detection of reduced plasmalogen (C16-C18 fatty acid) abundance at birth, followed by accumulation of phytanic acid within the first year (Braverman *et al.,* 2001; Irving *et al*., 2008; Krakow *et al*., 2003). Genetic sequencing identifies the specific lesion affecting *PEX7* and confirms the diagnosis. Disease severity correlates with variant functionality; the most severe RCDP1 symptoms are due to failed import of PTS2 client enzymes, indicating loss of *PEX7* function (Braverman *et al*., 1997; Braverman *et* al., 2001). In flies, an increase in total non-esterified fatty acid (NEFA) abundance was detected in homozygous *Pex7* mutant third instar larvae (Di Cara *et al*., 2019) suggesting an analogous role for fly *Pex7*. RCDP1 symptoms that manifest later in life include congenital cataracts, mild to severe mental retardation and recurrent respiratory tract infections due to neurologic compromise (Braverman *et al.,* 2001). Human *PEX7* thus has an important role in nervous system development and homeostasis. This study revealed fly *Pex7* has consistent proximity to the embryonic nervous system (Figure 4), peak expression and CNS cortex transcript localization during stage 17 (Figures 4 and 5), and embryonic sub-lethality when its expression is altered in neuroblasts (Figures 6-8) which together strongly imply neuro-specific functional conservation (Campos-Ortega and Hartenstein, 1985; Martinez Arias, 1993). The similarity in fly/human Pex7 DNA/protein sequences and predicted domains (Figure 9) support our inference, as does a recent report that *D. melanogaster* Pex7 can restore mis-localized Thiolase, a canonical PTS2-bearing enzyme, to peroxisomes in fibroblasts from RCDP1 patients (Di Cara *et al*., 2019).

**Figure 9.**
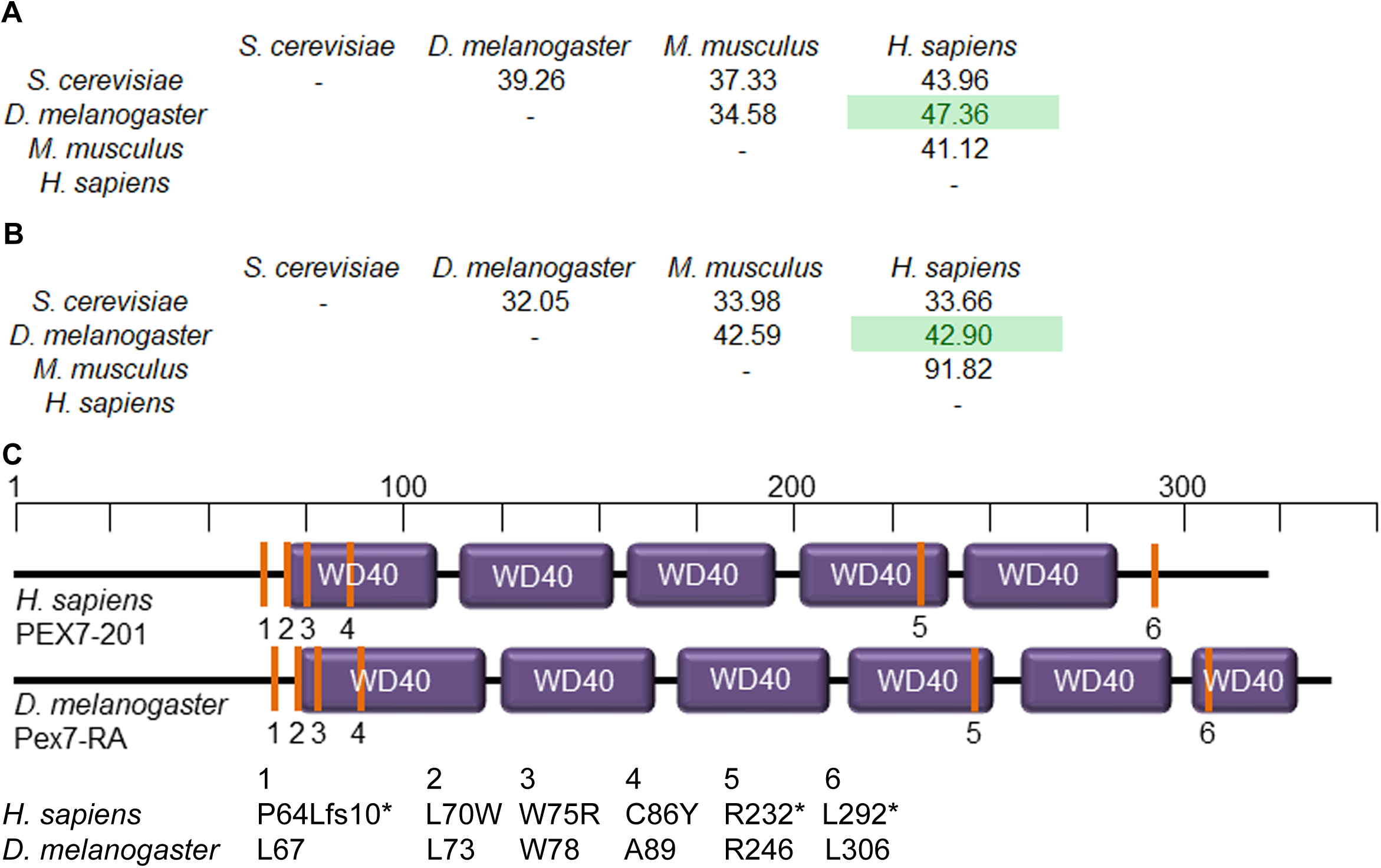
*Pex7* sequence and domain identity in select eukaryotes. (A) Primary isoform coding sequence (CDS) percent identity matrix (PIM) revealed fly/human CDS identity is higher than that of mouse/human. (B) Primary protein isoform PIM revealed fly/human protein sequence identity is within 5 % that of their DNA sequences however it is less than half that of mouse/human. (C) Human and fruit fly protein domain comparison. Labeled are human disease-associated residues and alignment-predicted *D. melanogaster* equivalents (orange bars), with the specific lesions detailed below.

It appears that *Pex7* is necessary for embryonic CNS development and its expression is regulated in a cell-lineage specific manner. Given that the canonical PTS2 pathway may be absent from *Drosophila* (Faust *et al.,* 2012; Baron *et al*., 2016) *Pex7* is likely not a PTS2 receptor protein. To contrast, *Pex7* mutations do cause systemic changes in lipid levels (Di Cara *et al.,* 2019), and Pex7 is targeted to peroxisomes in some cell types (Baron *et al.,* 2016). This suggests a role for Pex7 in some aspect of peroxisome function. We propose *D. melanogaster* Pex7 regulates lipid species key to CNS development, e.g. ether lipids or other phospholipid species.

## Author Contributions

Experiments were designed by AJS and CP and conducted by CP. Data analysis was performed by CP.

## Acknowledgements

We gratefully acknowledge the contributions of Richard A. Rachubinski, John Locke, Matthew Anderson-Baron, Virginia Pimmett, Xiao Li, Francesca Di Cara, Julie Haskins, Sarah Hughes and Sophie Keegan.

